# Reproducible transcriptional modules define glioblastoma ecosystems across independent cohorts

**DOI:** 10.64898/2026.05.20.726700

**Authors:** Heewon Seo

## Abstract

Glioblastoma (GBM) comprises a complex ecosystem of malignant, immune, vascular and neural transcriptional states. However, it remains difficult to determine which gene expression programmes are reproducibly recovered across independent cohorts and profiling platforms, because programme-level analyses are sensitive to cohort composition, technical context and factorization rank. Here, we analyzed three public GBM datasets—GLASS and IVYGAP bulk RNA-seq cohorts and the HEILAND Visium spatial transcriptomics cohort—to examine whether deconvolution-derived programmes could be organized into shared cross-cohort modules. Integrating 279 programmes inferred by consensus non-negative matrix factorization identified eight transcriptional communities, including myeloid immune microenvironment, neuronal and synaptic, oligodendrocyte and myelin, developmental, tumour-associated mesenchymal or hypoxic, proliferative, and ciliated or ependymal-like modules, as well as one cohort-restricted community. Community activity showed coherent associations with independent annotations: the myeloid community correlated with ESTIMATE immune score and inversely with tumour purity; the oligodendrocyte and myelin community was reduced in recurrent tumours; and ciliated or ependymal-like and neuronal communities showed modest exploratory associations with overall survival. Spatial projection onto Visium data provided qualitative support for the histological coherence of several modules, while also highlighting the limits of spot-level interpretation. Together, these results provide a proof-of-concept that cross-cohort integration can recover recurrent transcriptional structure across heterogeneous GBM datasets and offer an interpretable framework for comparing gene expression programmes while preserving cohort-specific signal and uncertainty in biological assignment.

## Introduction

Glioblastoma (GBM) is the most common and aggressive primary brain tumour in adults and is characterized by profound intra- and intertumoural heterogeneity^1^. Single-cell and spatial transcriptomic studies have shown that GBM comprises co-existing transcriptional programmes spanning malignant cell states and the surrounding microenvironment^2^, including mesenchymal-like, classical-like and proneural-like tumour programmes^3^, together with microglial, macrophage, oligodendrocyte and neuronal signals contributed by infiltrating immune cells^4^ and adjacent brain parenchyma^5^. These studies have reframed GBM as a complex ecosystem rather than a uniform tumour entity. However, it remains unclear which transcriptional programmes are consistently recovered across independent cohorts, tissue contexts and profiling platforms.

Matrix factorization approaches, including non-negative matrix factorization (NMF)^6,7,8^ and consensus NMF (cNMF)^9^, are widely used to resolve such structure by decomposing transcriptomic data into interpretable gene expression programmes and their sample-level activities. These methods can recover biologically meaningful latent factors, but the inferred programmes are often sensitive to the factorization rank, cohort composition, and technical context. As a result, programmes derived in one study are not readily comparable to those identified in another, particularly when datasets differ in platform, anatomical sampling, tissue composition or preprocessing. Existing efforts to align programmes across studies have often relied on pairwise heuristics, such as overlap among highly weighted genes or correlation to reference signatures, but these approaches do not provide a general framework for identifying higher-order transcriptional structure shared across datasets.

We previously developed the Spatial Omics Toolkit, which introduced a data-driven strategy for rank selection and metagene network analysis in spatial transcriptomics and enabled systematic evaluation of gene expression programme structure in a neurodegenerative disease context^10^. Here, we extend that analytical framework to the increasingly common setting in which multiple GBM cohorts, generated using distinct platforms and study designs, are available for joint analysis. Using this approach, which we term *sotk2*, we integrated cohort-derived programmes within a correlation network and used community structure to distinguish recurrent cross-cohort modules from cohort-restricted signal.

Applying this framework to three public GBM datasets—GLASS, IVYGAP and HEILAND—we identified reproducible transcriptional modules that capture major components of the GBM ecosystem across bulk and spatial transcriptomic data. These modules corresponded to tumour-intrinsic, immune, neural and developmental programmes and showed distinct associations with independent molecular, clinical and spatial annotations, including molecular subtype, anatomical compartment, recurrence status, immune infiltration, tumour purity, overall survival and histological context. Together, these results define a reproducible cross-cohort transcriptional architecture for GBM and provide a general strategy for comparing gene expression programmes across heterogeneous datasets.

## Results

### Cross-cohort integration identifies recurrent transcriptional modules across glioblastoma datasets

To assess whether gene expression programmes (GEPs) inferred independently from distinct GBM cohorts could be integrated into a shared transcriptional framework, we analyzed cNMF-derived programmes from three public datasets spanning different assay types, tissue contexts and levels of resolution (Fig. 1A). Concatenation of basis matrices across all included ranks and cohorts yielded 279 GEPs defined across 26,316 shared genes, placing all programmes in a common gene space for direct comparison (Fig. 1B). Pairwise Spearman correlations across the 279 GEPs revealed substantial positive structure both within and between cohorts. As expected, within-cohort correlations were generally stronger, reflecting shared assay conditions and preprocessing. However, positive cross-cohort correlations were also common, indicating that at least a subset of pro grammes recovered independently from bulk RNA-seq and spatial transcriptomic data capture related biological structure.

**Figure 1.**
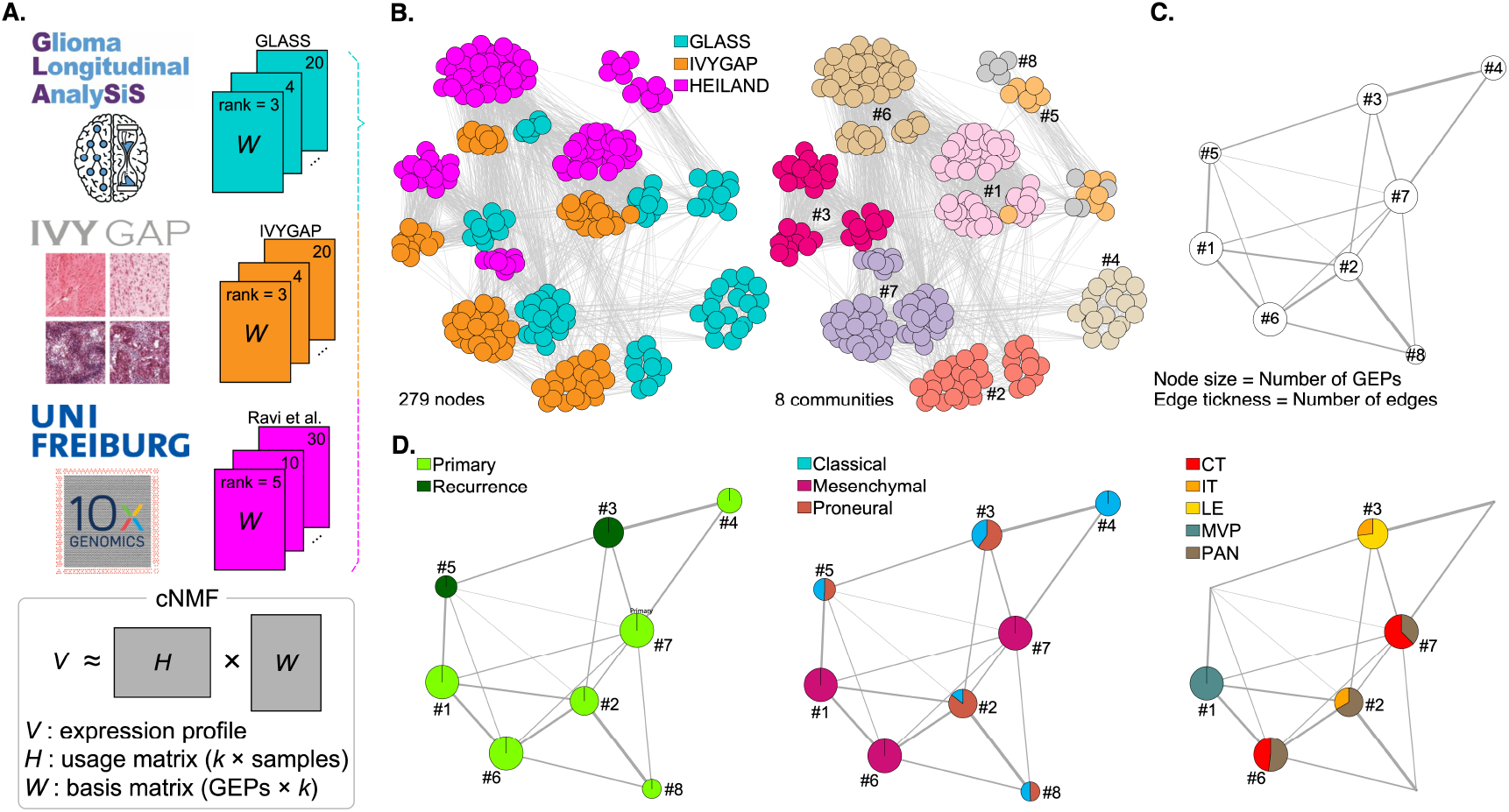
Conceptual framework and cross-cohort integration of multiple GBM datasets using *sotk2*. **(A)** Three cohorts with distinct platforms and resolutions were deconvolved using consensus non-negative matrix factorization to derive gene expression programmes (GEPs). Program (*W*) matrices were used to compute cross-program correlations and to construct a GEP correlation network. **(B)** GEP-level correlation networks. Left: applying a correlation threshold of >0.3 retained 279 GEPs and positive correlations were represented as edges; node colors indicate cohort. Right: community detection (*fast greedy*) partitioned the network into eight communities; node colors indicate community membership. **(C)** Community-level abstraction of the GEP network, where each node represents a community and edges summarize inter-community connectivity. Node size is proportional to the number of GEPs assigned to the community, and edge thickness is proportional to the number of inter-community edges in the underlying GEP network. **(D)** Community-level networks with sample-annotation overlays shown as pie charts based on standardized chi-squared residuals (enrichment/depletion relative to expectation). Left: GLASS primary versus recurrence. Center: GLASS molecular subtype (Verhaak signatures). Right: IVYGAP anatomical features (CT, cellular tumour; IT, infiltrating tumour; LE, tumour’s leading edge; MVP, microvascular proliferations; PAN, palisading cells around necrosis).

We therefore constructed a thresholded network based on positive correlations (ρ > 0.3), retaining all 279 GEPs as connected nodes. Community detection identified eight communities (Fig. 1C). Community sizes ranged from 9 GEPs (Community #8) to 59 GEPs (Community #6), indicating variability in the extent to which distinct programmes recurred across cohorts. Importantly, most communities spanned multiple factorization ranks, suggesting that they were not driven solely by a single decomposition setting. A community-level abstracted network provided a compact summary of relationships among modules and served as a scaffold for downstream annotation (Fig. 1D). These analyses provide an initial proof of concept that independently inferred GEPs from heterogeneous GBM datasets can be organized into higher-order cross-cohort modules, while also retaining cohort-restricted structure where present.

### Community-level gene signatures identify distinct tumour, neural and microenvironment-associated modules

To assign provisional biological identities to the recovered communities, we identified metagene-associated genes (MAGs) for GLASS-derived GEPs and then extracted genes recurrently represented across GEPs within each module. This procedure yielded interpretable gene signatures for seven of the eight communities, spanning immune, neural, developmental and tumour-associated programmes (Fig. 2; Table 1). Given the use of public bulk and spatial transcriptomic data, these assignments should be viewed as exploratory biological interpretations rather than definitive cell-state definitions.

**Table 1.**
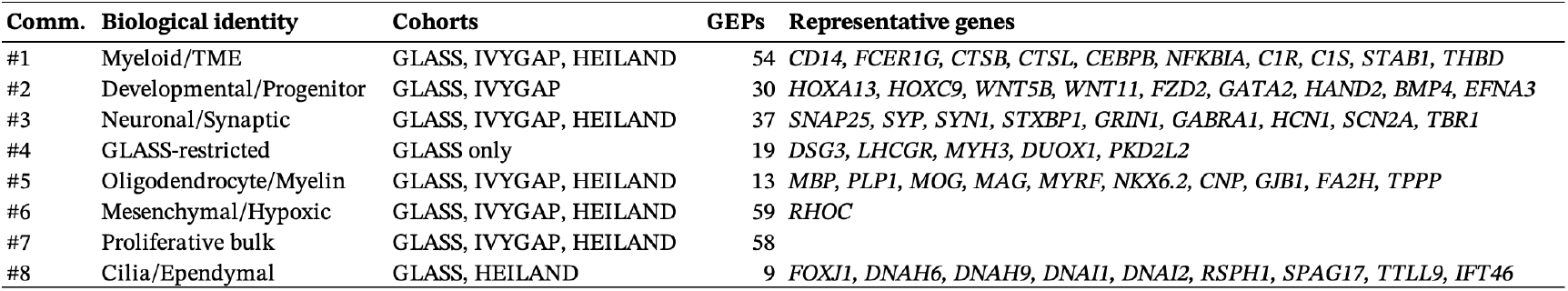
Summary of eight GBM community modules identified by *sotk2*.

**Figure 2.**
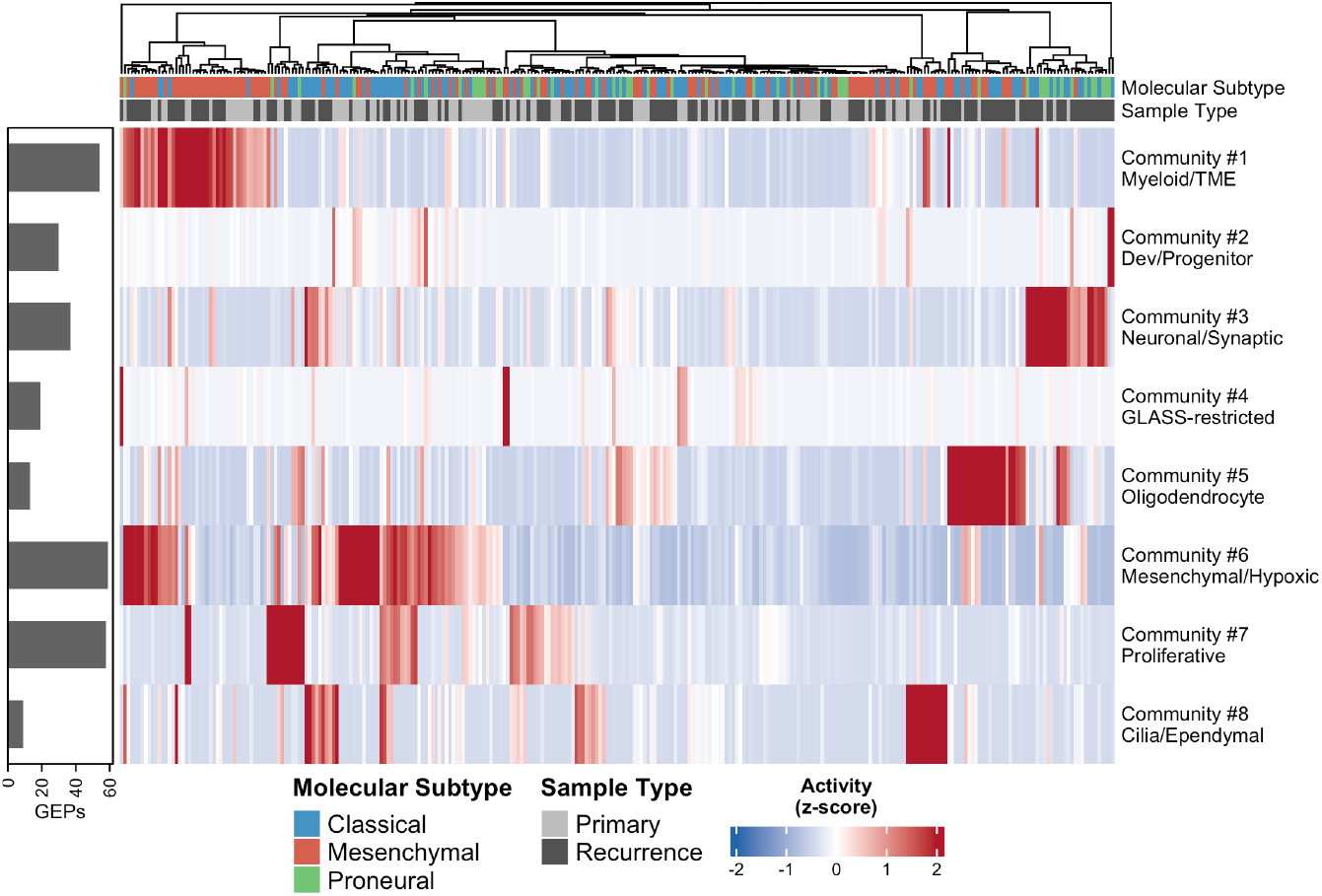
Community activity across GLASS samples reveals biologically distinct transcriptional modules. Heatmap of community activity scores for all 291 GLASS samples (columns) across eight GEP communities (rows). Activity scores represent the geometric mean of normalized GEP usage across all GEPs assigned to each community, computed independently per sample. Values are z-scored per community (row) prior to display. Samples are hierarchically clustered (Euclidean distance, Ward.D2 linkage); community order is fixed (Community #1–#8). Top annotation bar indicates molecular subtype (Classical, blue; Mesenchymal, red; Proneural, green) and sample type (Primary, light grey; Recurrence, dark grey). Left bar indicates the number of GEPs assigned to each community. Community biological identities are based on metagene-associated gene (MAG) analysis and contributing community gene extraction.

#### Community #1: myeloid and immune microenvironment-associated programme

Community #1 comprised 54 GEPs distributed across all three cohorts, making it one of the most broadly shared modules in the network. Its contributing genes included canonical myeloid and immune-associated markers such as *CD14, FCER1G, FCGRT, LAPTM5, HCST, LILRB3* and *OSCAR*, together with complement components (*C1R, C1S*), inflammatory regulators (*CEBPB, NFKBIA*) and lysosomal genes (*CTSB, CTSC, CTSL, CTSZ*). This pattern is consistent with a myeloid-rich tumour microenvironment programme, likely reflecting contributions from macrophages and microglia, although the present analysis does not resolve their relative abundance or state.

#### Community #2: developmental or progenitor-associated programme

Community #2 comprised 30 GEPs shared between GLASS and IVYGAP, but absent from HEILAND. Its contributing genes included multiple HOX family members, WNT pathway components (*WNT5B, WNT6, WNT9A, WNT11, FZD2, FZD9, FZD10*), and developmental transcription factors, including *GATA2, HAND2, PAX2, LHX1, ISL2, TBX1, TBX3*, and *PITX1*. These features support the interpretation that it is a developmental or progenitor-associated programme. Its absence from the HEILAND cohort may reflect limited detectability at Visium spot resolution, differences in tissue sampling, or cohort-specific representation.

#### Community #3: neuronal and synaptic programme

Community #3 comprised 37 GEPs across all three cohorts and showed one of the strongest and most coherent gene signatures in the analysis. Contributing genes included synaptic vesicle and neurotransmission components such as *SNAP25, SYP, SYN1, SYT1, STXBP1, RAB3A* and *NSF*, together with neurotransmitter receptors and ion channels including *GRIN1, GABRA1, GABRB2, HCN1, SCN2A* and *KCNA4*. This pattern is consistent with neuronal and synaptic signals, most plausibly reflecting non-malignant neural tissue captured in or adjacent to tumour specimens, rather than a canonical tumour-intrinsic GBM programme. However, because these data are derived from bulk and spot-level profiles, this interpretation remains inferential.

#### Community #4: GLASS-restricted cohort-specific signal

Community #4 comprised 19 GEPs, all from GLASS, and lacked a clear biological signature. Its contributing genes did not resolve into an obvious GBM-relevant programme, suggesting that this module may represent cohort-specific technical structure, sample-specific variation or low-confidence biological signal. We therefore retained it as a cohort-restricted community but did not interpret it further biologically.

#### Community #5: oligodendrocyte and myelin-associated programme

Community #5 was the smallest cross-cohort module (13 GEPs), but displayed one of the most specific gene signatures. Contributing genes included canonical oligodendrocyte and myelin markers such as *MBP, PLP1, MOG, MAG, MOBP, OPALIN, CNP, MYRF* and *NKX6*.*2*. This pattern is consistent with oligodendrocyte and white-matter-associated signal. As with the neuronal module, the present data do not distinguish whether this reflects non-malignant tissue admixture, regional sampling differences, or more complex relationships between the tumour and the surrounding brain.

#### Community #6: mesenchymal or hypoxia-associated tumour programme

Community #6 was the largest module in the network (59 GEPs) and was dominated by HEILAND-derived programmes. It yielded relatively few genes shared across the majority of member GEPs, indicating substantial internal heterogeneity. Nevertheless, its behaviour across external annotations was consistent with a broad mesenchymal- or hypoxia-associated tumour programme. Because this community is not strongly anchored by a compact shared gene set, its biological interpretation should be considered provisional.

#### Community #7: proliferative tumour-associated programme

Community #7 comprised 58 GEPs and, like Community 6, showed broad internal heterogeneity and limited convergence on a concise shared gene signature. Its strongest anatomical enrichment was in cellular tumour regions, supporting interpretation as a proliferative or tumour-bulk-associated programme. However, given the limited gene-level specificity, this assignment should also be regarded as exploratory.

#### Community #8: ciliated or ependymal-associated programme

Community #8 comprised 9 GEPs from GLASS and HEILAND and showed a highly coherent motile cilia signature. Contributing genes included *FOXJ1*, multiple axonemal dyneins (*DNAH6, DNAH9, DNAH12, DNAI1, DNAI2*) and additional ciliary structure genes such as *RSPH1, RSPH4A, SPAG1, SPAG17, TTLL9* and *IFT46*. This pattern is consistent with a ciliated or ependymal-associated signal. As above, the most likely interpretation is a contribution from non-malignant tissue context, although the present analysis does not directly establish anatomical origin.

Together, these signatures suggest that cross-cohort programme integration can recover biologically interpretable modules spanning tumour-intrinsic and non-malignant components of the GBM ecosystem, while also preserving uncertainty where the evidence is weaker.

### Community activity is coherently organized across molecular subtype and anatomical compartment

We next asked whether community activity was associated with independent molecular and anatomical annotations not used for network construction. These analyses were intended as orthogonal validation of community structure, while remaining exploratory given the retrospective use of public datasets. In the GLASS cohort, Community #1 showed the highest activity in mesenchymal tumours (Fig. 3A), consistent with prior observations that mesenchymal GBM is enriched for immune and inflammatory signatures. Community #3 showed higher activity in classical and proneural tumours than in mesenchymal tumours, in keeping with its neuronal interpretation. Community #6 was also the highest in mesenchymal samples, whereas Community #7 was more evenly distributed across molecular subtypes, consistent with a more broadly represented tumour-associated programme. In IVYGAP, which samples anatomically distinct histological compartments, community activity also showed structured spatial associations (Fig. 3B). Community #1 was highest in microvascular proliferations, consistent with a vascular and immune-rich microenvironment. Community #3 showed the strongest activity at the leading edge and within the infiltrating tumour, supporting an association with the tumour–brain interface. Community #6 was most active in palisading cells around necrosis, consistent with a hypoxia-associated programme. Community #5 showed higher activity in cellular tumour and infiltrating tumour, which may reflect co-sampling of white-matter-associated signal in these regions. Taken together, the concordance between GLASS subtype associations and IVYGAP compartment enrichments supports the view that the recovered communities capture biologically meaningful structure across independent cohorts. At the same time, these associations should be interpreted as supportive rather than definitive, because they derive from indirect annotation rather than direct experimental validation.

**Figure 3.**
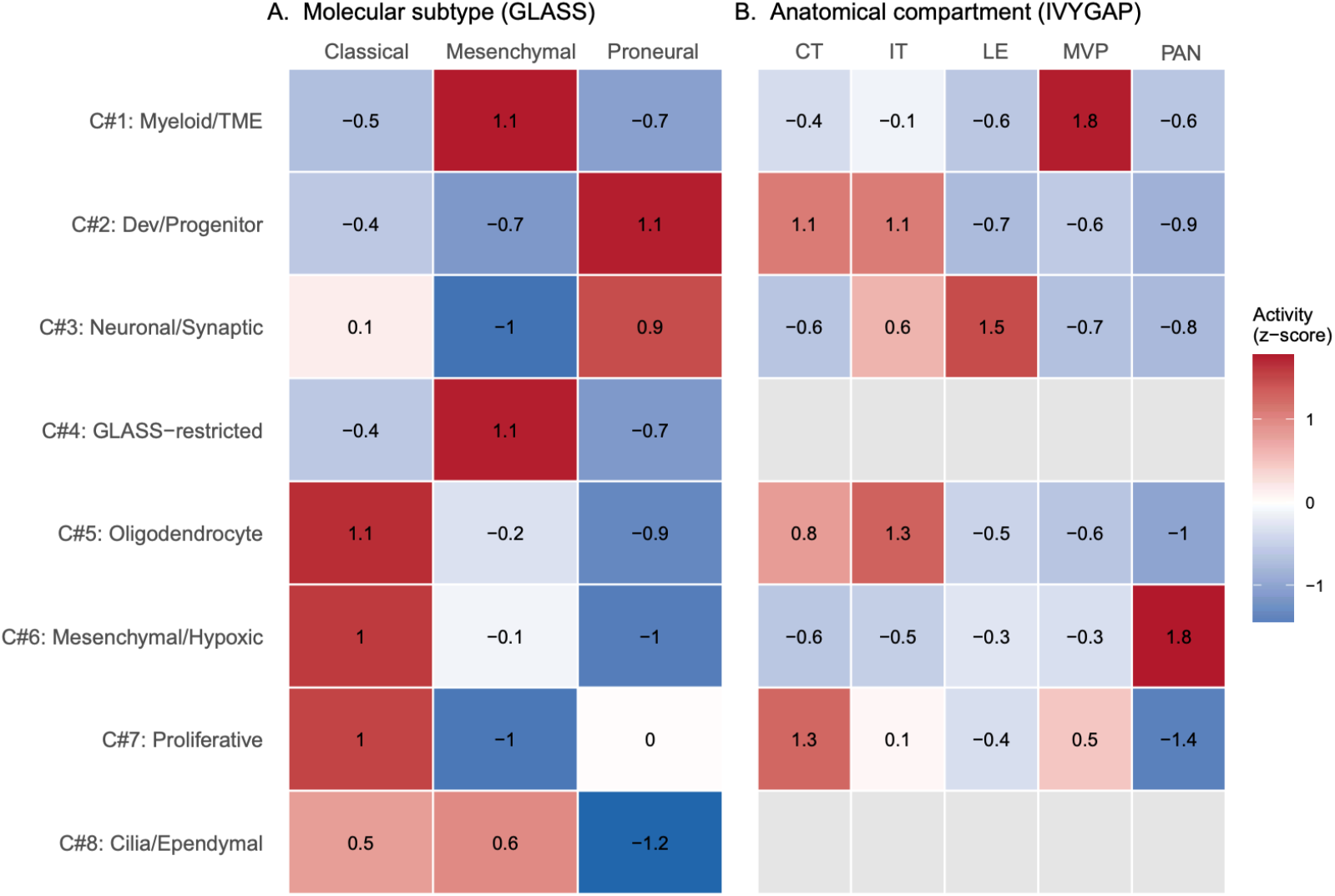
Community activity is coherently organized with molecular subtype and anatomical tumour compartment. Tile heatmap showing mean community activity z-scores across independent annotation categories. Values are z-scored per community (row) to enable comparison across communities with different absolute activity ranges; z-score values are printed in each cell. **(A)** Mean community activity by GLASS molecular subtype (Classical, Mesenchymal, Proneural; Verhaak classification). **(B)** Mean community activity by IVYGAP anatomical compartment: cellular tumour (CT), infiltrating tumour (IT), leading edge (LE), microvascular proliferations (MVP), and palisading cells around necrosis (PAN). Community biological identities are defined in Table 1.

### Longitudinal analysis suggests depletion of neural white-matter-associated signal and increased tumour programme activity at recurrence

The longitudinal structure of the GLASS cohort enabled an exploratory comparison between primary (*N* = 154) and recurrent (*N* = 137) GBM samples. Several communities showed significant differences in activity between these groups (Fig. 4A). The strongest shift was observed for Community #5, which was markedly reduced at recurrence (Wilcoxon *P* < 0.0001). Given its oligodendrocyte and myelin-associated signature, this result suggests a relative loss of white-matter-associated signal in recurrent tumours. Because these data derive from publicly available bulk RNAseq data and do not directly measure cell composition or treatment effects, we interpret this finding conservatively as a longitudinal association rather than as evidence of a specific biological mechanism. Community #3 was also lower at recurrence (*P* = 0.013), suggesting reduced neuronal-associated signal in recurrent samples. This could reflect differences in tissue context, biopsy location or changes in the relative contribution of tumour and surrounding brain, but these possibilities cannot be distinguished in the present data. By contrast, both Community #6 (*P* = 0.003) and Community #7 (*P* = 0.002) showed higher activity at recurrence. This pattern is consistent with greater representation of tumour-associated programmes in recurrent samples, although the broad and heterogeneous nature of these communities limits more specific interpretation. No significant differences were observed for Communities #1, #4 or #8. Overall, these results suggest that recurrence in GBM is accompanied by shifts in the balance between tumour-associated and non-tumour-associated transcriptional signals, but the current analysis should be regarded as exploratory and hypothesis-generating.

**Figure 4.**
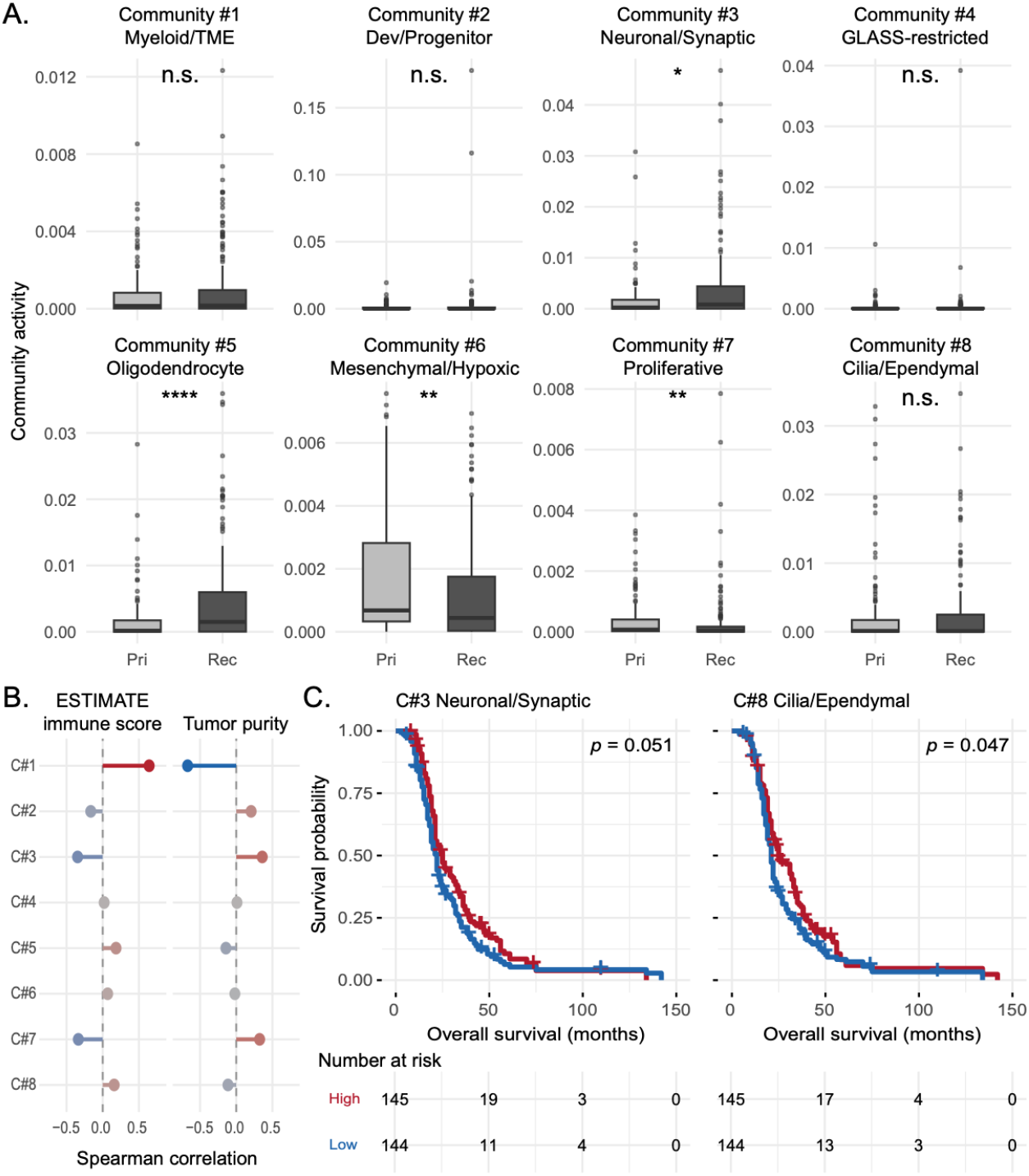
Community activity dynamics across disease progression, tumour microenvironment composition, and patient survival. (**A)** Boxplots of community activity scores in primary (light grey, *N*=154) and recurrent (dark grey, *N*=137) GLASS samples across all eight communities. Wilcoxon rank-sum test significance: **** *p* < 0.0001, ** *p* < 0.01, * *p* < 0.05, n.s. *p* >= 0.05. **(B)** Lollipop plots showing Spearman correlation coefficients between community activity and ESTIMATE immune score (left) and tumour purity (right) across GLASS samples. Positive coefficient (red) indicates co-directional association; negative coefficient (blue) indicates inverse association. **(C)** Kaplan-Meier overall survival curves for Community #3 (log-rank *p* = 0.051) and Community #8 (*p* = 0.047). Samples were dichotomized at the median community activity into high (red) and low (blue) groups. Survival was compared using the log-rank (Mantel-Cox) test (*N*=289 with survival data). At-risk tables are shown below each curve.

### Myeloid community activity tracks immune infiltration and inversely correlates with tumour purity

To further assess community identity, we correlated GLASS community activity scores with two independent computational annotations: the ESTIMATE immune score and tumour purity estimates. Community #1 showed the strongest association with either variable, correlating positively with immune score (Spearman *r* = 0.64) and negatively with tumour purity (*r* = −0.67) (Fig. 4B). These findings provide orthogonal support for interpreting Community #1 as an immune-and myeloid-associated programme. Community #3 showed the opposite pattern, with a lower immune score and higher tumour purity associated with stronger neuronal activity. This is consistent with the possibility that the neuronal signal is more prominent in samples with less immune infiltration, although the relationship remains correlative. Communities #6 and #7 both showed a moderate positive correlation with tumour purity, consistent with their tentative assignment as tumour-associated modules. These associations indicate that the recovered communities capture major compositional axes in public GBM transcriptomic data, particularly the contrast between immune-rich and tumour-dominant samples.

### Ciliated and neuronal-associated communities show exploratory associations with overall survival

We next examined whether community activity was associated with overall survival in GLASS (*N* = 289 with survival data). For each community, samples were dichotomized at the median activity score and compared using the log-rank test (Fig. 4C). Given the modest effect sizes, the use of public observational data and the multiple communities tested, these analyses should be regarded as exploratory. Community #8 showed an association with longer overall survival: patients with high activity had a median survival of 24.0 months, compared with 21.0 months in the low-activity group (log-rank *P* = 0.047). Community #3 showed a similar directional trend that did not meet conventional significance (*P* = 0.051). No other community showed a significant association with survival in this univariable analysis. Because these associations are modest and unadjusted for potential confounders, we do not interpret them as evidence of independent prognostic value. Instead, they suggest that neural and ciliated tissue-context-associated programmes may warrant further evaluation in future survival-analysis studies.

### Spatial projection provides a limited but supportive in situ context for recovered modules

To complement the bulk analyses with spatial information, we projected community assignments onto Visium v1 spots from a representative HEILAND section and compared them with available histological segmentation labels (Fig. 5). Interpretation of these results requires caution because cNMF was performed at the spot level rather than the sample level. Each Visium spot can capture transcripts from multiple neighbouring cells and may therefore reflect mixed tissue composition rather than a single discrete histological state. Spatial correspondences were therefore interpreted as qualitative support for community identity rather than as strict one-to-one validation. With these limitations in mind, several spatial patterns were consistent with the bulk-derived community annotations. One Visium community was enriched for white matter and showed little representation in cellular tumour regions, consistent with the oligodendrocyte- and myelin-associated programme identified in bulk data. Another was concentrated in infiltrative regions, in keeping with the neuronal and synaptic module associated with the tumour–brain interface. More spatially distributed communities aligned with cellular tumour or vascular-hyperplastic regions, broadly consistent with tumour-associated and immune-associated modules identified in the integrated network. Thus, although the spot-level Visium analysis does not permit strict one-to-one mapping between histological labels and inferred communities, it provides supportive spatial context for the broader biological interpretation of the recovered modules. In combination with gene-level signatures and cohort-level annotations, these results suggest that the integrated communities reflect spatially structured features of GBM tissue organization.

**Figure 5.**
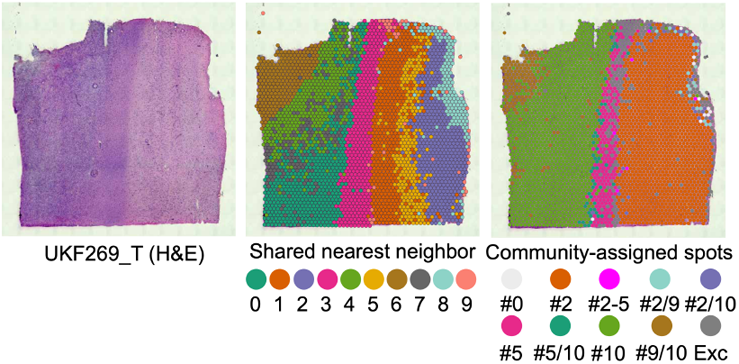
Spatial projection of *sotk2* community assignments onto Visium histology corroborates bulk-derived community biology. (left) H&E image of the UKF269_T glioblastoma section. (center) Seurat shared nearest neighbor clusters overlaid on Visium spots. (right) *sotk2* community labels projected onto spots; spots may map to multiple communities. Hashmarks (e.g., “#2”) indicate the community identifier (here, Community #2); “#0” denotes spots with memberships spanning more than two communities, and “Exc” denotes spots that could not be assigned under the selected criteria.

### Cross-cohort integration defines an exploratory transcriptional framework for glioblastoma ecosystems

Across three independent public datasets, cross-cohort integration of deconvolution-derived programmes recovered recurrent transcriptional modules corresponding to immune, neural, oligodendrocyte, developmental and tumour-associated signals. Several of these modules were supported by consistent associations with molecular subtype, anatomical compartment, recurrence status, immune infiltration, tumour purity and spatial histology. At the same time, some communities—particularly Communities #6 and #7—remained biologically broad and incompletely resolved, underscoring the limits of the present data and the need for further refinement. Taken together, these results establish a proof of concept that cross-cohort integration can recover a reproducible, though still exploratory, transcriptional framework for describing GBM ecosystems across heterogeneous datasets.

## Discussion

We applied *sotk2* to integrate GEPs inferred independently from three public GBM cohorts spanning bulk RNA-seq and spatial transcriptomics and recovered eight community modules that summarize recurrent transcriptional structure across datasets. Most communities were represented in more than one cohort, suggesting that at least part of the programme-level organization of GBM is reproducible across distinct study designs, tissue contexts and profiling platforms. In this setting, the principal value of the analysis is not to definitively establish new cell states, but to provide a cross-cohort framework for identifying shared versus cohort-restricted transcriptional signals in heterogeneous public data.

A central challenge in comparing deconvolution-derived programmes across studies is that inferred factors depend on rank choice, cohort composition, assay modality and preprocessing. By representing cohort-derived programmes in a common correlation network and identifying higher-order communities, *sotk2* provides a way to compare programme structure without requiring direct batch correction, reference mapping or one-to-one matching of factors across datasets. In the present study, this strategy recovered modules with interpretable associations to immune, neural, oligodendrocyte, developmental and tumour-associated signals, while also preserving a cohort-restricted community lacking clear biological coherence. This distinction between shared and restricted structure is important in practice because public datasets often contain both reproducible biology and study-specific signals that should not be forced into a unified interpretation. The biological annotations assigned to individual communities should, however, be regarded as provisional. They are based on recurrent gene signatures and orthogonal associations with available metadata, rather than on direct experimental validation. This is particularly relevant for modules inferred from bulk RNA-seq, where observed programmes may reflect differences in tissue admixture, sampling location, tumour purity or microenvironmental composition rather than discrete cellular states. The same caution applies to spot-level spatial transcriptomic data, in which each Visium spot can capture transcripts from multiple neighbouring cells. Accordingly, the labels used here— such as myeloid, neuronal, oligodendrocyte, mesenchymal or ciliated—are best understood as working descriptors of dominant signal composition rather than definitive assignments of cellular origin.

This conservative interpretation is especially important for Communities #6 and #7. Both were large and recurrent across datasets and showed structured associations with external annotations, but neither resolved into a compact, highly specific shared gene signature. These communities, therefore, appear to capture broad tumour-associated structure rather than sharply delimited biological states. In this sense, the present analysis is informative but incomplete: it suggests that cross-cohort integration can recover stable higher-order tumour modules, while also showing that some components of GBM transcriptional heterogeneity remain difficult to resolve cleanly using public bulk and spot-level data alone. The survival analyses should likewise be interpreted cautiously. The associations observed for the neuronal- and cilia-associated communities were modest in magnitude and derived from univariable stratification of a retrospective public cohort. These findings are therefore best treated as exploratory signals rather than evidence of independent prognostic value. More generally, this study was not designed to support causal or clinically actionable inference. Instead, it provides a proof of concept that community-level programme organization can be linked to external molecular and clinical annotations, potentially guiding future hypothesis-driven studies. The spatial analyses also require careful qualification. Community assignments in HEILAND were derived from spot-level factorization rather than from whole-sample profiles, and individual spots may contain mixed cellular and histological compositions.

In summary, this study establishes a proof of concept for cross-cohort integration of deconvolution-derived transcriptional programmes in GBM. Across three public datasets, the analysis recovered recurrent modules with consistent, though exploratory, associations to molecular subtype, anatomical compartment, recurrence status, immune infiltration, tumour purity and spatial tissue context. At the same time, the results underscore the limits of biological interpretation from public bulk and spot-level data and highlight the need for follow-up studies with matched orthogonal validation. Framed in this way, *sotk2* provides a practical and interpretable approach for organizing transcriptional programme structure across heterogeneous cohorts while preserving uncertainty where the evidence remains incomplete.

## Materials and Methods

### Public glioblastoma cohorts

We analyzed three public GBM transcriptomic datasets:

1. The GLASS dataset comprises longitudinal bulk RNA-seq profiles from primary and recurrent GBM tumours (*N* = 291 samples; 154 primary and 137 recurrent)^11,12^, together with molecular subtype annotations according to the Verhaak classification (classical, mesenchymal and proneural)^3^, ESTIMATE immune and stromal scores^13^, tumour-purity estimates, and overall survival data (*N* = 289 with survival annotation).
2. The IVYGAP dataset comprises bulk RNA-seq profiles from anatomically distinct GBM regions (*N* = 270 samples), annotated by histological compartment as cellular tumour (CT), infiltrating tumour (IT), leading edge (LE), microvascular proliferation (MVP) and palisading cells around necrosis (PAN)^14^.
3. The HEILAND dataset comprises Visium v1 spatial transcriptomic profiles from 27 GBM tissue sections (70,929 spots in total), with accompanying histological segmentation labels (Cellular, Infiltrative, Necrosis, Necrotic_Edge, Vascular_Hyper, White_Matter and Grey_Matter)^15^ and Seurat shared nearest neighbour (SNN) cluster assignments computed using Seurat^16^.

### Consensus non-negative matrix factorization

GEPs were inferred independently for each cohort using consensus non-negative matrix factorization (cNMF; v1.4)^7^. The following filtering and regularization parameters were used: `removeAbove = 10, removeBelow = 0, alpha = 1`, and `density threshold = 2` . For GLASS and IVYGAP, ranks *k* = 2–60 were evaluated. For HEILAND, which showed greater sparsity at the Visium spot level, a coarser rank grid (*k* = 5, 10, 15, 20, 25, 30) was used. All remaining cNMF parameters were left at package defaults. Analyses were run with a fixed random seed (`seed = 14`).

### Cross-cohort programme integration

To integrate inferred programmes across cohorts, we applied *sotk2* to cNMF outputs from GLASS, IVYGAP and HEILAND. For GLASS and IVYGAP, programmes from ranks k = 3–10, 15 and 20 were retained, whereas for HEILAND, programmes from ranks k = 5, 10, 15, 20, 25 and 30 were included. GEPs were then represented as nodes in a correlation network, and higher-order communities of related programmes were identified using a network-based integration strategy conceptually related to that described by Hofree et al. (2013)^17^. Basis matrices (*W*) from all selected ranks and cohorts were concatenated, yielding 279 GEPs across 26,316 genes. Pairwise Spearman correlations were then calculated between all GEPs using pairwise-complete observations. A correlation threshold of 0.3 was applied to construct a GEP correlation network, retaining only positive correlations. Community detection was performed on this network using the fast greedy algorithm^18^ with a fixed seed (`seed = 1234, niter = 1000`). Community-weighted and cohort-weighted layouts `(commWeight = 100, cohortWeight = 10`) were used for visualization only and did not affect community assignment.

### Metagene-associated genes

MAGs were identified for each GEP in the GLASS cohort using the `getMAGs()` function in *sotk2*, which implements an approach adapted from Tsukahara et al^19^. First, the top 200 genes ranked by *W*-matrix loading were selected for each GEP. Second, among these genes, those with a Spearman correlation greater than 0.2 between per-sample gene expression and the corresponding H-matrix usage were retained. Community-level contributing genes were then identified using `contributingCommunityGenes()` with a proportion threshold of 0.5, such that genes were retained if they appeared among the MAGs of more than 50% of GEPs within a given community.

### Community activity scores

For each sample in GLASS and IVYGAP, a community activity score was calculated for each community as the geometric mean of normalized persample GEP usage values across all GEPs assigned to that community. To improve comparability across ranks, GEP usage values were row-normalized within each rank before aggregation.

### Association with molecular and clinical annotations

Community activity scores were linked to sample-level annotations in GLASS, including the Verhaak molecular subtype, sample type (primary versus recurrence), the ESTIMATE immune score, and tumour purity. Differences in community activity between sample groups were assessed using the Wilcoxon rank-sum test. Associations between community activity and continuous variables, including immune score and tumour purity, were evaluated using Spearman correlation. For IVYGAP, anatomical annotation was derived from sample identifiers that encoded histological compartments (CT, IT, LE, MVP, and PAN).

### Survival analysis

Overall survival data from GLASS were linked to sample-level community activity scores using case barcodes. For each community, samples were dichotomized at the median activity score, and overall survival was compared between high- and low-activity groups using the log-rank (Mantel–Cox) test. Median overall survival was calculated for each group.

### Spatial analysis

For spatial validation, community assignments from one representative HEILAND Visium section (UKF269_T) were projected onto spot-level histological segmentation labels provided in the HEILAND metadata. Spot-to-community assignments were derived from a precomputed sotk2 object generated with the Shiny application’s default parameters. Co-occurrence between community assignments and histological segmentation labels was summarized in a contingency table.

## Software and code availability

sotk2 is distributed as an open-source R package: https://github.com/Snyder-Institute/sotk2. Precomputed cNMF outputs and the bundled sotk2 object used in this analysis are deposited on Zenodo (https://doi.org/10.5281/zenodo.18063318). End-to-end analysis scripts are provided under “inst/scripts/” in the package repository.

## Acknowledgements

We gratefully acknowledge support from the Snyder Institute for Chronic Diseases at the University of Calgary.

## Competing interest statement

The author declares no competing interests.

## Notes

### Competing Interest Statement

The authors have declared no competing interest.

